# A partially averaged system to model neuron responses to interferential current stimulation

**DOI:** 10.1101/2022.05.23.493095

**Authors:** Eduardo Cerpa, Matías Courdurier, Esteban Hernández, Leonel E. Medina, Esteban Paduro

## Abstract

The interferential current (IFC) therapy is a noninvasive electrical neurostimulation technique intended to activate deep neurons using surface electrodes. In IFC, two independent kilohertz-frequency currents purportedly intersect where an interference field is generated. However, the effects of IFC on neurons within and outside the interference field are not completely understood, and it is unclear whether this technique can reliable activate deep target neurons without side effects. In recent years, realistic computational models of IFC have been introduced to quantify the effects of IFC on brain cells, but they are often complex and computationally costly. Here, we introduce a simplified model of IFC based on the FitzHugh-Nagumo (FHN) model of a neuron. By considering a modified averaging method, we obtain a non-autonomous approximated system, with explicit representation of relevant IFC parameters. For this approximated system we determine conditions under which it reliably approximates the complete FHN system under IFC stimulation, and we mathematically prove its ability to predict nonspiking states. In addition, we perform numerical simulations that show that the interference effect is observed only for a narrow set of IFC parameters and, in particular, for a beat frequency no higher than about 100 [Hz]. Our novel model tailored to the IFC technique contributes to the understanding of neurostimulation modalities using this type of signals, and can have implications in the design of noninvasive electrical stimulation therapies.

## 1 Introduction

### 1.1 Motivation

Electrical stimulation therapies are used to treat the symptoms of a variety of nervous system disorders and diseases, including Parkinson’s disease, epilepsy, and chronic pain, among others [13]. In these therapies, electrical currents intended to modulate the activity of nerve fibers or neurons are delivered to the tissues through implanted or surface electrodes, commonly in the form of a train of pulses with a repetition rate of a few tens of Hz. The interferential current (IFC) therapy [16] is a transcutaneous electrical stimulation technique in which two pairs of skin electrodes are placed diagonally opposed such that the paths of two independently generated kilohertz-frequency (KHF) currents produce an interference effect where they intersect. In IFC, sinusoidal currents of slightly different frequencies, *e.g*., 1000 and 1050 [Hz], are applied continuously at a fixed amplitude, and it is intended that neurons experience a fully modulated signal resulting from the arithmetic summation of the two sinusoids, *i.e*., the interference effect [23, 15] (Figure 1). In IFC, the actual stimulation signal that reaches neurons is not known, *i.e*., neurons can experience a purely KHF sinusoid (unmodulated signal) or a signal with certain degree of modulation depending on the location within the interference field [1]. Volume conductor models may be used to determine the spatiotemporal distribution of currents for IFC, and coupled to model of neurons, can be used to predict neuron responses to modulated and unmodulated signals [14, 6]. In fact, recent computational modeling studies have analyzed the effects of IFC in the brain using realistic 3D models of the head and/or morphologically-detailed biophysical models of neurons [3, 10, 9, 21, 2]. However, there is still a lack of fundamental understanding of the mechanisms of neuron activation under IFC stimulation, perhaps in part due to the scarce availability of analytical approaches that provide immediate insight. The goal of this study is to develop a simplified model of IFC that is both manageable and with explicit representation of the interference effect on neurons.

**Figure 1:**
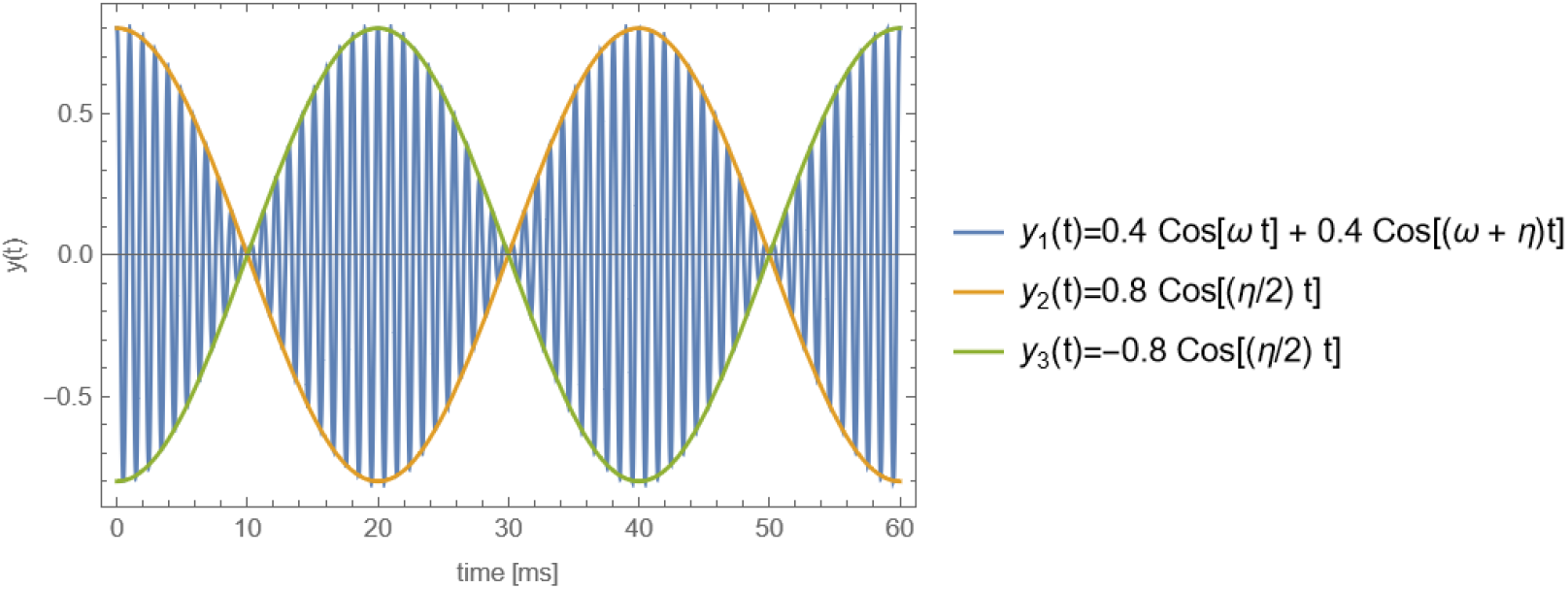
Interference pattern of two oscillatory terms with frequencies *ω*/(2π) = 1000 [Hz] and (*ω* + *η*)/(2π) = 1050 [Hz], and with amplitudes of *A* = 0.4 and *B* = 0.4 during a time window of 60 [ms]. The arithmetic addition of these two sinusoids results in a modulated waveform with a beat frequency of 1050 – 1000 = 50 [Hz]

### 1.2 Model description

We consider the FitzHugh-Nagumo (FHN) model of a neuron [4], which is a generalization of the Van der Pol oscillator. This model captures key features of the Hodgkin and Huxley (HH) [8] equations of neuron membrane dynamics, but with lower dimension. The FHN model has been widely used in the study of chaotic systems because of the presence of the phenomena of relaxation oscillations and the existence of canard solutions [5], [11]. The FHN system is described by the following equations

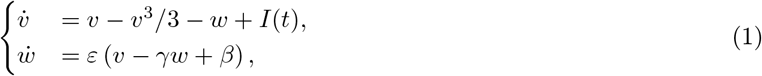

where *ε* > 0 (typically small), *γ* > 0, 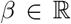 and *I*(*t*) is an input current. Similar to the HH model [4, 8], *v* represents the voltage across the cell membrane, and *w* is a restitution variable analog to the ion channels in the HH model.

In this study, we focused in the case for which the system has a single stable equilibrium point. More precisely, the parameters *β*, *γ*, *ε* are chosen so that the system

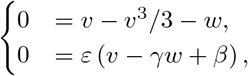

has a unique solution 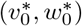 and additionally it satisfies 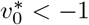. To make this conditions explicit, we use that 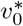 is defined as the solution of the equation *f*(*v*) = *v*^3^ – 3(1 – 1/*γ*)*v* + 3*β*/*γ* = 0. For a depressed cubic, the condition for uniqueness of the real root can be written as

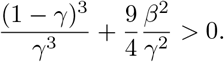

Additionally, to have 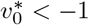, since *f* (*v*) → ∞ as *v* → ∞, it is enough that *f* (–1) > 0. Thus,

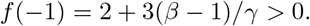

This gives the following conditions on the parameters:

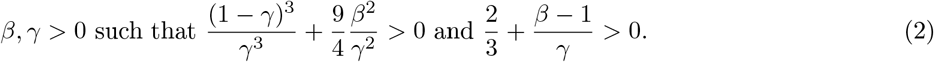

In previous work [24, 17], the FHN model was studied using excitation inputs of the form

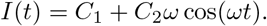

In these studies, the averaging method was used to deal with the source term and obtain autonomous systems, which can be studied with dynamical systems methods. See [20] for an excellent presentation of the averaging method in different contexts.

In this paper, we present a novel model of IFC stimulation based on the FHN formulation that incorporates two KHF sinusoidal currents with slightly different frequency as input signals

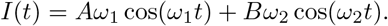

Our goal is to determine if, by exploring these two frequency components, it is possible to obtain a new response of the system. We use partial averaging of the equation to introduce an approximation of the problem via a non-autonomous ODE. Further, since the interference field purportedly affecting neurons comprises a low frequency envelope resulting from the arithmetic addition of two sinusoids, *i.e*. the beat frequency, we aim at making explicit the effect of such envelope. Since the approximated system is non-autonomous, its analysis poses some challenges, as we will see later.

### 1.3 The approximated model

Consider the FHN model (1) with an input current of the form

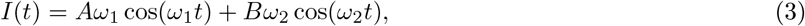

where, *ω*_1_ ≫ 1, *ω*_2_ = *ω*_1_ + *η* for some small beat frequency *η* > 0. The magnitude of the coefficients *A* and *B* can be related to the strength and/or the distance to the current source. For example, the potential field, *V_e_*, generated by a current point-source of amplitude *I*, is 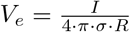, with *R* the distance to the source.

#### Definition 1.1

(Partially Averaged System). *Let* 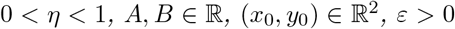 *and β, γ satisfying* (2). *We say that* (*V, W*) *is a solution of the Partially Averaged System if it satisfies the following non-autonomous equations*

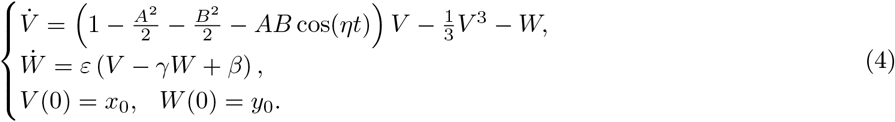

The derivation of this approximation is presented in Section 2. We note that for *A* = 0 or *B* = 0, *i.e*., no interferential currents, we recover the approximated system obtained using the averaging method [24]. Numerical simulations of the Partially Averaged System are presented in Section 5.

### 1.4 Relaxation oscillations and action potentials

In excitable cells, an action potential or spike is a rapid rise and subsequent fall of the voltage across the cell membrane generated by the flux of ions [8]. This definition is useful in a different context, but we need a more precise mathematical definition for our analysis. From the point of view of chaotic systems, relaxation oscillators like the FHN neuron have the property that solutions are very unstable near a curve known as the separatrix. The separatrix determines a boundary for regions in the domain where the system exhibits different behaviors, and for neuron models we can use this change of behavior as a definition of action potential. Consequently, we call action potentials to the solutions of the system that cross this separatrix, solutions also known as relaxation oscillations [22].

For FHN, we know that the separatrix can be described as the solution of the FHN system that intersects the null line 0 = *v* – *v*^3^/3 – *w* at its local maximum, which for this cubic equation corresponds to the point (1, 2/3). The separatrix in Figure 2 is found by solving the system backwards starting at the point (1, 2/3). An important property about the separatrix in the stable case, is that it always lies to the left side of the vertical line *v* = 1 and therefore any solution (*v, w*) that does not cross the line *v* = 1 has no action potentials. This criteria will be used in Section 5 to justify that if *v*(*t*) ≤ 1 then no action potential was elicited. We note that a more realistic range for the membrane potential, *e.g*. ~100 [mV] as measured experimentally [8], can be obtained by simply incorporating appropriate offset parameters in the FHN system. In this case, a similar separatrix analysis can be performed to determine thresholds for action potential identification.

**Figure 2:**
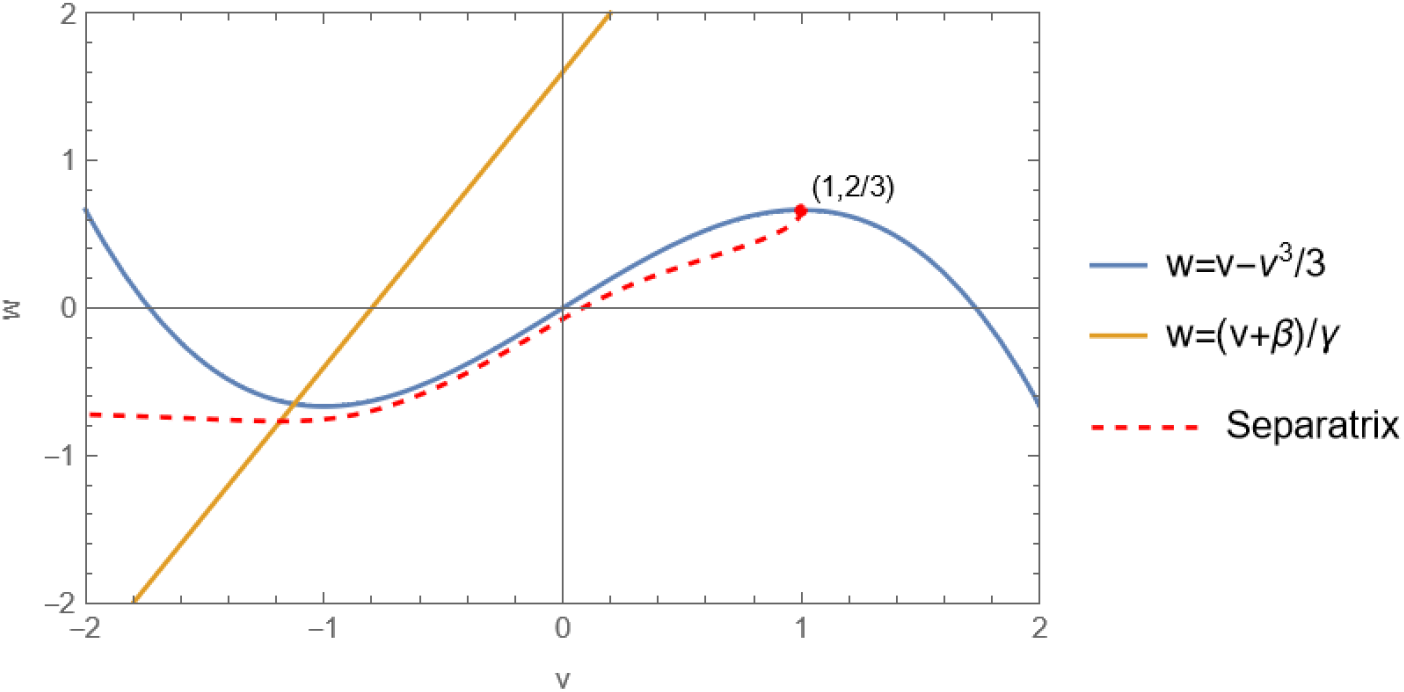
Characteristics of the FHN system. The null lines, *w* = *v* – *v*^3^/3 and *w* = (*v* + *β*)/*γ*, and the separatrix of the FHN system are shown in the (*v, w*) plane. In this case, *β* = 0.8, *γ* = 0.5, *ε* = 0.08

### 1.5 Main Results

In this work we present and characterize the properties of the Partially Averaged System (4) that describes neuron behavior under IFC stimulation. We also illustrate how that system (4) can be used to predict properties of the more complex system (1) under stimulations as signals (3). For this purpose, we study the dependence of the solution on the parameters *A, B* and *η* and determine the regions for which the Partially Averaged System (4) accurately predicts the behavior of the FHN system (1).

The first result in this study is stated in Theorem 1, which specifies the conditions for *A, B* under which the absence of relaxation oscillations in the Partially Averaged System (4) can predict the absence of relaxation oscillations in the full FHN system (1). Next, in Theorems 2 and 3 we present certain properties of the Partially Averaged System (4) that explain some of the inherent difficulties for neuron activation via IFC stimulation. This is done by proving that:

i. For small amplitudes of stimulation, *i.e*., small values of parameters *A, B*, the Partially Averaged System (4) cannot generate action potentials.
ii. For large beat frequency *η* the Partially Averaged System (4) cannot generate action potentials.

Taken together, these results show that the region of parameters *A, B* and *η* where IFC stimulation may activate neurons is rather small and therefore the parameters require a careful and well understood calibration. For a precise formulation of our results, the following definition is useful.

#### Definition 1.2.

*Let* 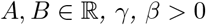 *satisfying* (2). *The numbers v*_0_, *w*_0_ *are defined as the unique real valued solution of*

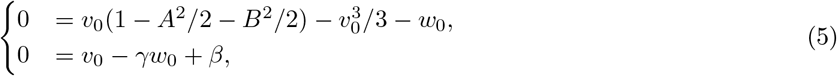

*additionally because of condition* (2) *it is easy to show that v*_0_ < –1.

**Remark 1.1.** *If A* ≠ 0 *or B* ≠ 0, *then* (*v*_0_, *w*_0_) *is not an equilibrium point for the Partially Averaged System* (4). *However, as we will see in the proof, it is a convenient choice to study the system around this point*.

Our first result is about the approximation of system (1) by the Partially Averaged System (4).

#### Theorem 1

(Approximation result). *Fix ε, γ, β* > 0 *satisfying* (2) *and consider the solution* (*v, w*) *of the FHN system* (1) *under the IFC stimulation*

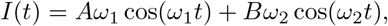

*where ω*_1_ ≫ 1, *ω*_2_ = *ω*_1_+*η*, |*η*| ≤ 1, *A*, 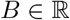, *and with initial condition* 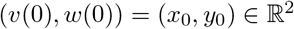. *Additionally, let* (*V, W*) *be the solution of the Partially Averaged System* (4) *with initial condition* (*V* (0), *W*(0)) = (*x*_0_, *y*_0_). *Then, there exists c*_0_ > 0, *M* = *M*(*ε, γ, β*) > 0, *ω*_0_ = *ω*_0_(*ε, γ, β*) *and some* 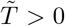 *independent of ω*_1_ *such that if* max{|*A*|, |*B*|} ≤ *M*, *ω*_1_ ≥ *ω*_0_ *and*

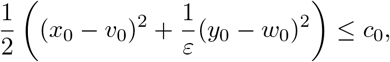

*where* (*v*_0_, *w*_0_) *is given by* (5)*. Then we have the estimate*

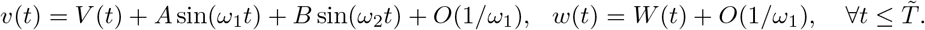

Next, we study the stability of the Partially Averaged System (4) in the following result.

#### Theorem 2

(Stability of the Partially Averaged System). *Let* 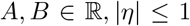, *fix ε, γ, β* > 0 *satisfying* (2), *let* (*v*_0_, *w*_0_) = (*v*_0_(*A, B*), *w*_0_(*A, B*)) *be given by* (5) *and consider the solution* (*V, W*) *of system* (4) *with initial condition* 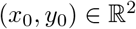.

a. *There exists a non-empty set* 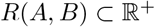 *such that if the initial condition* 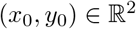 *satisfies*

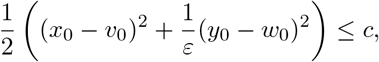

*for c* ∈ *R*(*A, B*), *then we have the estimate*

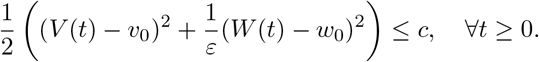
b. *Suppose that A, B and v*_0_ *satisfy*

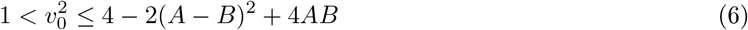

*then there exists constants c*_1_ > 0 *and* 0 < *c*_2_ < *c*_3_ *such that*

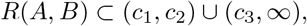

*where the constants c*_1_, *c*_2_, *c*_3_ *satisfy*

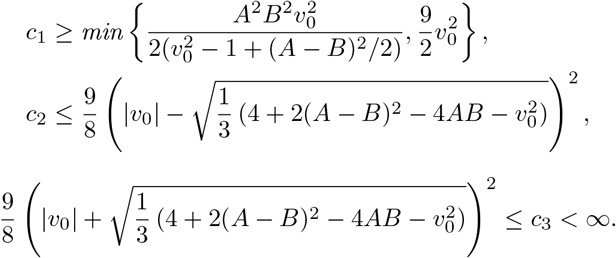 *Additionally, there exists K*_0_ = *K*_0_(*ε*, *β*, *γ*) > 0 *such that if the parameters A, B*, *γ, ε and v*_0_ *satisfy*

i. max{|*A*|, |*B*|} ≤ *K*_0_,
ii. 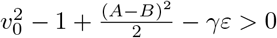,

*then the intersection R*(*A, B*) ⋂ (*c*_1_, *c*_2_) *is non empty*.
c. *Suppose that A, B*, *γ, ε and v*_0_(*A, B*) *satisfy* (6) *and assumptions i. and ii. as in item (b). Given δ* > 0 *there exists M* = *M*(*ε*, *γ*, *β*, *δ*) > 0 *such that if* max{|*A*|, |*B*|} ≤ *M, then there exists* 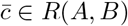, *such that if the initial condition* 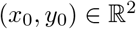 *satisfies*

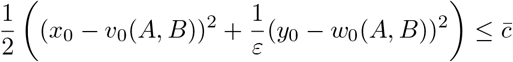

*then the solution* (*V, W*) *of system* (4) *with initial data* (*x*_0_, *y*_0_) *satisfies*

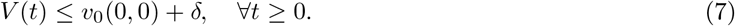

The main idea used on the proof of Theorem 2 is to study the existence of invariant ellipses for system (4). This can be seen as an analogous to the technique of invariant rectangles [18] used to prove the global existence of the FHN model in a cable, with the additional difficulty that because of the IFC stimulus, the invariant regions cannot be arbitrarily small.

The previous theorem studies the role of the current amplitudes *A* and *B*, whereas the following result is related to the role of the beat frequency *η*. We note that a key feature of IFC stimulation is that the beat frequency *η* is relatively small.

#### Theorem 3

(No action potential for large beat frequency). *Let* 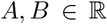, *fix ε*, *γ*, *β* > 0 *satisfying* (2) *and* (*v*_0_, *w*_0_) *be given by* (5). *There exists η*_0_ > 0 *such that for all η* ≥ *η*_0_ *the solutions* (*V, W*) *of* (4) *with initial conditions satisfying*

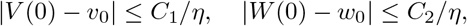

*are such that*

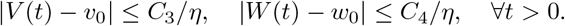

*for some constants C*_1_, *C*_2_, *C*_3_, *C*_4_ > 0.

The rest of the paper is organized as follow. In Section 2 we derive the Partially Averaged System (4) as a simplified model of IFC stimulation based on the FitzHugh-Nagumo (FHN) model of a neuron. In Section 3 we analyze the Partially Averaged System (4) and present the proofs of Theorem 2 and Theorem 3. Section 4 is devoted to determine conditions under which the Partially Averaged System reliably approximates the complete FHN system under IFC stimulation. This section contains the proof of Theorem 1. Numerical simulations and conclusions are presented in Section 5 and Section 6, respectively. Finally, some intermediate technical results are included in the Appendix A.

## 2 Derivation of the Partially Averaged System

Consider the FHN model (1) with a IFC input current (3), i.e.,

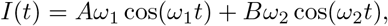

where, *ω*_1_ ≫ 1, *ω*_2_ = *ω*_1_ + *η* for some small beat frequency *η* > 0. We note that the size of the coefficients *A* and *B* can be related to the strength of the stimulation signal and the distance from the neuron to the source. Motivated by the analysis of a single high frequency input [24, 17] we look for solutions of the form

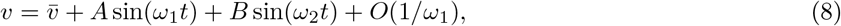

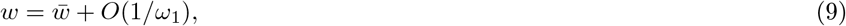

where 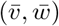 represent some kind of averaging. To look for the equations for 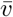 and 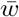 we neglect the error terms and substitute 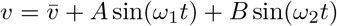 and 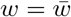 in the first equation in (1). We obtain

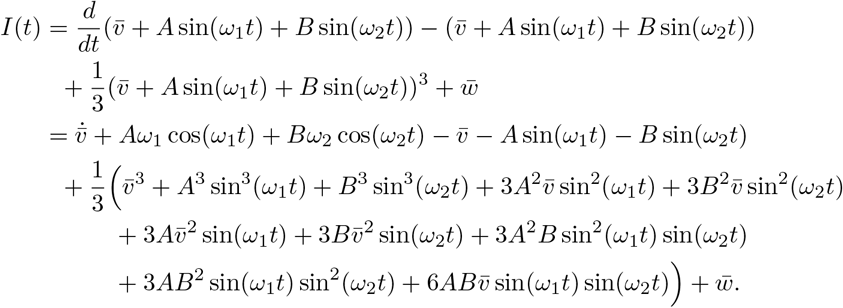

As the term *Aω*_1_ cos(*ω*_1_*t*) + *Bω*_2_ cos(*ω*_2_*t*) can be canceled out with *I*(*t*), we get

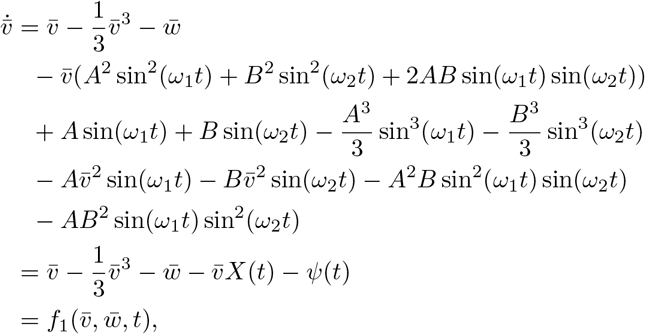

where we denote

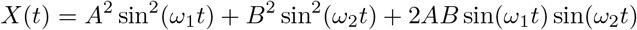

and

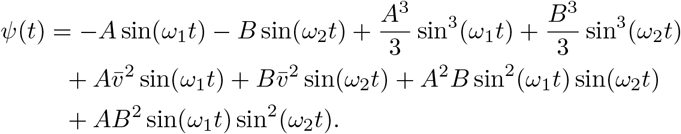

Here, we are assuming that 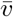 and 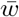 vary very slowly compared to the oscillatory terms. Under this assumption, it is reasonable to consider the following averaging

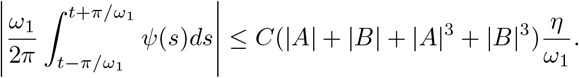

In *X*(*t*) we use the identities 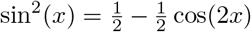 and 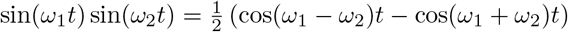 to get the following decomposition

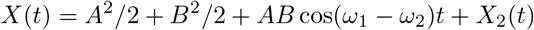

where 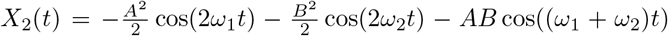. Taking the same averaging as before and using *ω*_1_ – *ω*_2_ = –*η* we get

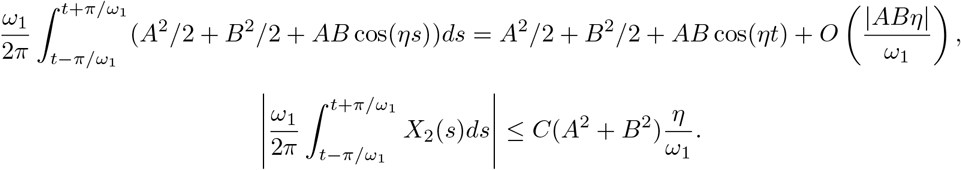

Now we look at the equation for *w* in (1). We can write

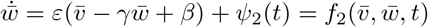

where *ψ*_2_ = *ε*(*A* sin(*ω*_1_*t*) + *B* sin(*ω*_2_*t*)). Putting all together we get from the averaging on [*t* – π/*ω*_1_, *t* + π/*ω*_1_]

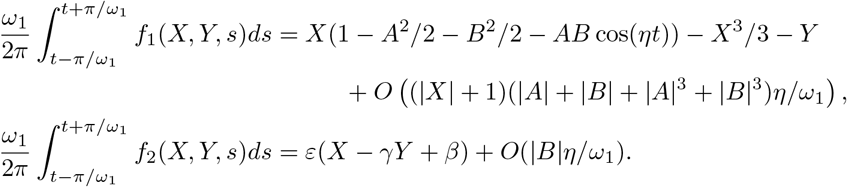

Therefore, by ignoring all the *O*(1/*ω*_1_) terms, we get the Partially Averaged System given by (4), i.e.,

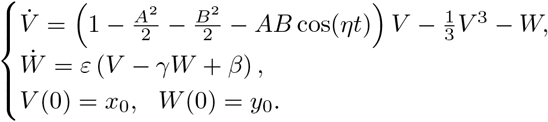

This system is non-autonomous, and has an explicit dependence on the beat frequency generated by the interference of the KHF input signals. We note that for *A* = 0 or *B* = 0, *i.e*., no interferential currents, we recover the approximated system obtained by the averaging method [24].

Later on in this paper, we will analyze the the extent to which this Partially Averaged System allows us to study properties of the FHN system (1) under IFC stimulations. In this context, the following result, whose proof can be found in the Appendix A, will be required.

### Proposition 4

(Error equation). *Let* 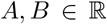, |*η*| ≤ 1, *and let ε*, *γ*, *β* > 0 *satisfying* (2). *Let* (*v, w*) *be the solution of the FHN system* (1) *and* (*V, W*) *be the solution of the Partially Averaged System* (4), *both with initial condition* (*v*_0_, *w*_0_) *given by* (5). *The functions*

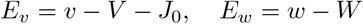

*are solutions of*

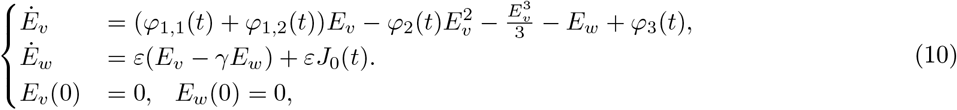

*where*

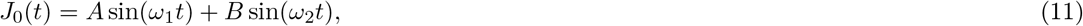

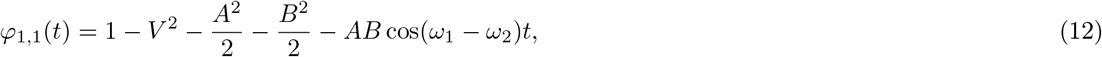

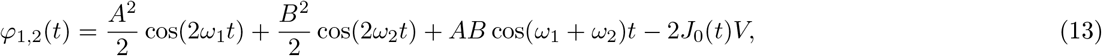

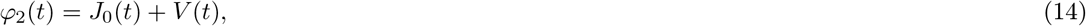

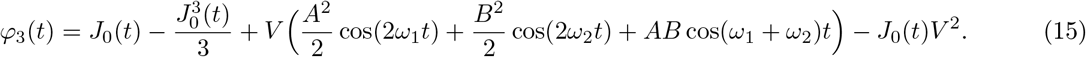

## 3 Analysis of the Partially Averaged System

The goal of this section is to establish properties of the solutions of the Partially Averaged System (4) under appropriate assumptions on the parameters of the equation. In particular, we want to know when the solutions stay near the point (*v*_0_, *w*_0_), given by (5).

### 3.1 Proof of Theorem 2

In this proof it is useful to rewrite the system (4) as follows

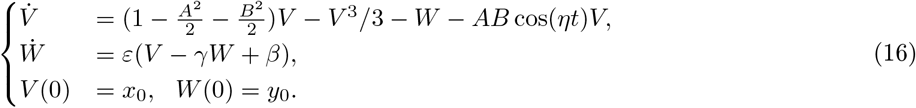

Next, using the change of variable *V* = *v*_0_ + *δv, W* = *w*_0_ + *δw* we can center the system (16) about the point (*v*_0_, *w*_0_) defined by (5), obtaining

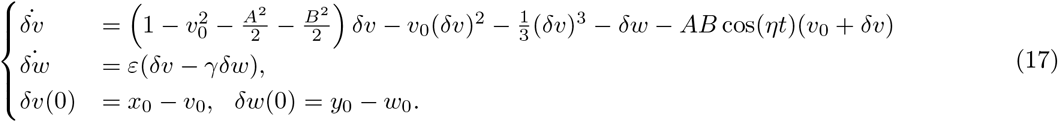

#### Proof of item (a)

We consider the nonlinear system (17) with initial conditions 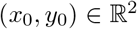. Our goal is to determine conditions for (*x*_0_, *y*_0_) such that the solution of this system is trapped in an ellipse. For this purpose, we look at the derivative of the quantity 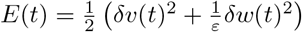, as follows

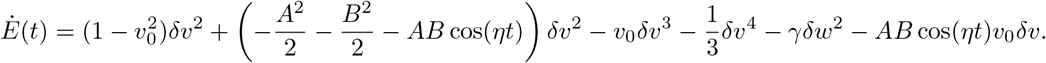

In virtue of that 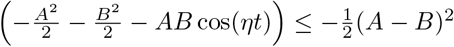, we define the set 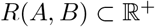 as

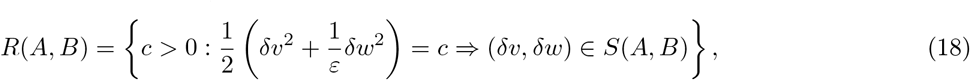

where the region 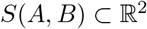 is defined by

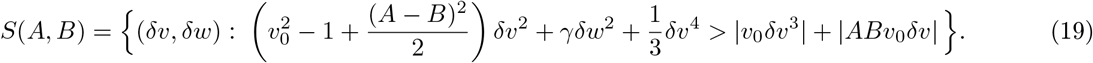

It is easy to see that if for some *t* > 0 we have (*δv*(*t*), *δw*(*t*)) ∈ *S*(*A, B*) then we can guarantee that *E*′(*t*) < 0. Finally, to conclude that the definition of the set *R*(*A, B*) suffices to guarantee that the solution is trapped, we use the following result whose proof can be found in the Appendix A.

##### Lemma 5.

*Let c* > 0 *and* 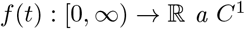 *function such that*

i. *f* (0) ≤ *c, and*
ii. *If f*(*t*) = *c for some t* ∈ (0, ∞), *then* 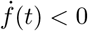.

*Then, f*(*t*) ≤ *c for all t* ∈ (0, ∞).

The last component to complete item (a) is to verify that the set *R*(*A, B*) is not empty. We claim that large ellipses are always completely contained in *S*(*A, B*). To demonstrate it, we look for *D*_1_ > 0 and *D*_2_ > 0 such that

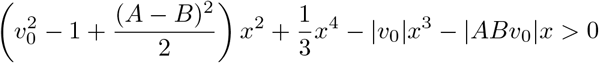

for *x* > *D*_1_ and

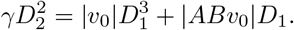

Thus, we can guarantee that

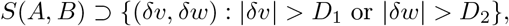

and therefore any ellipse outside of such rectangle is completely contained in *S*(*A, B*). We can take 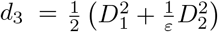 and conclude that (*d*_3_, ∞) ⊂ *R*(*A, B*). It follows that the set *R*(*A, B*) is non empty, which concludes the proof of item (a) of Theorem 2.

#### Proof of item (b)

We need to verify several assertions about the set *R*(*A, B*) defined by (18). This will be done in several steps:

Step 1) We show there is a region near the origin of forbidden values of *c* ∈ *R*(*A, B*).
Step 2) We show that, depending on the values of *A* and *B*, there exists an interval of forbidden values of *c* ∈ *R*(*A, B*).
Step 3) Finally, we show that under appropriate conditions we can ensure that the intersection (*c*_1_, *c*_2_) ⋂ *R*(*A, B*) is nonempty.

From the definition of the set *S*(*A, B*) (19) we can re-write the set *R*(*A, B*) as an intersection of a family of sets indexed by a parameter *α* in the following way

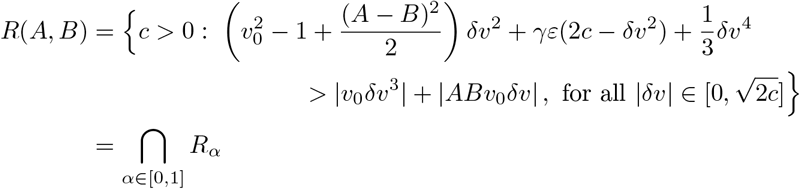

where

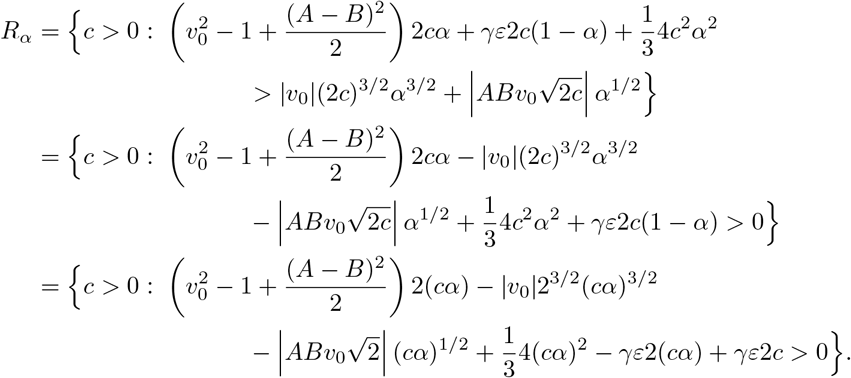

Since this set is defined by a family of conditions, we can get restrictions on the set *R*(*A, B*) by checking individual values for *δv* ∈ [0, 2c]. For α = 0 the condition becomes *R*_0_ = {*c* > 0 : *γε*2*c* > 0}, which is always satisfied for all *c* > 0. We get a more interesting condition when we check for *α* = 1, which is equivalent to the case 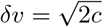.

##### Step 1) Lower bound for *c*_1_ and *c*_3_

To see that small ellipses do not satisfy the condition, we look at the 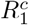

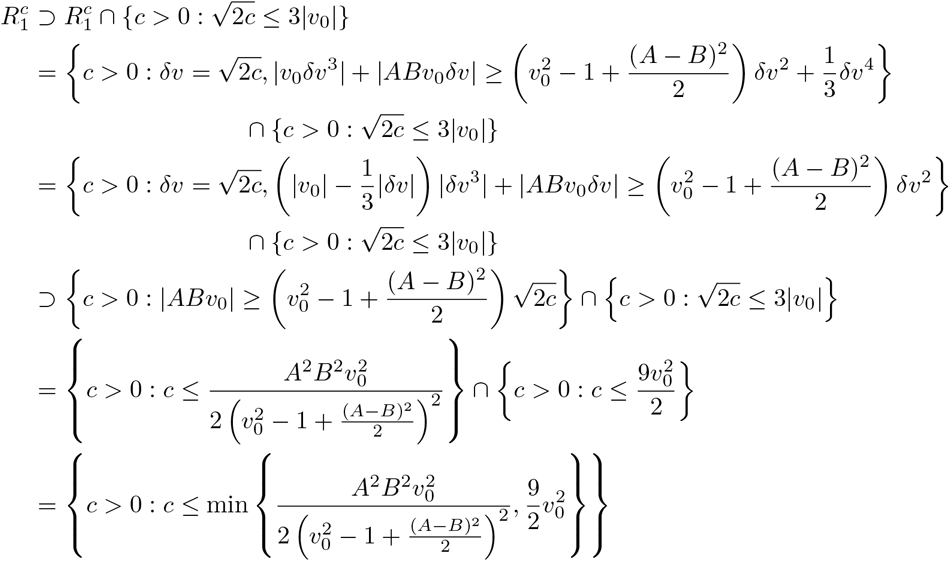

taking complements again we obtain the lower bound.

##### Step 2) Upper bound for *c*_2_ and lower bound for *c*_3_

The next bound is more delicate because we want to argue that depending on the values of *A* and *B* we can find a range of forbidden values for *c*. Intuitively, that corresponds to the region where the term *v*_0_*v*^3^ is dominant. To formalize this, we note that in the region where |*δv*| ≤ *ρ*|*v*_0_| for some 0 < *ρ* < 3, we can bound 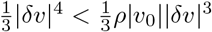 and therefore we have the inclusion

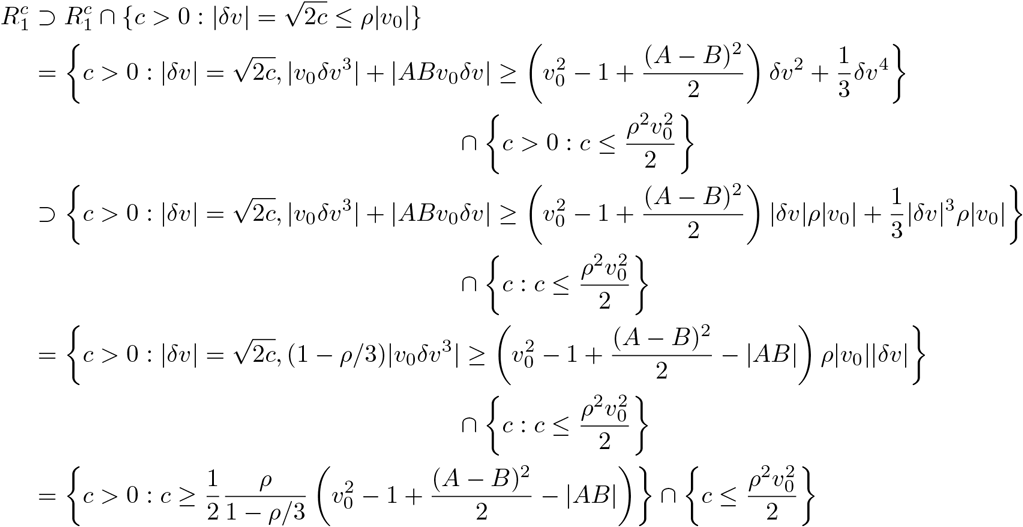

we get the following inclusion for 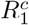

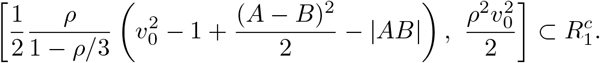

Since we know that for any value of *ρ* > 0 this represents a set that is contained in 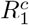, we want to know how big this union is. First, since the limits of the interval vary continuously with *ρ* > 0, we now that the union is also an interval. As *ρ* becomes too small or too big it is easy to see that the interval collapse and becomes empty. Therefore, we can find the limits of this union by looking at the values of *ρ* such that

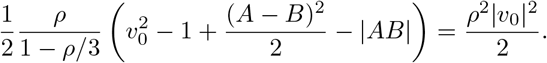

This can be solved easily using the quadratic formula

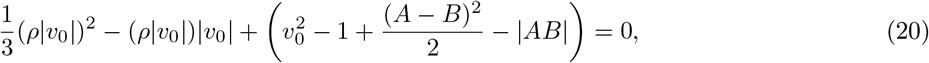

to get the solutions

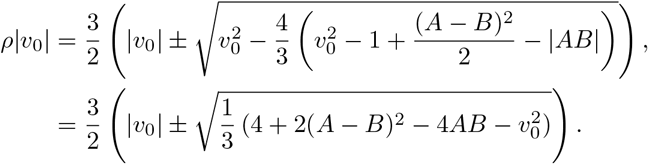

Notice that because of assumption (6) both solutions are real, additionally because there are two sign changes in (20) we know that both roots as strictly positive. We obtain the inclusion

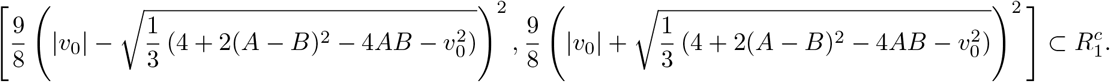

By taking complements and using Step 1) we conclude that

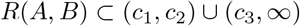

where

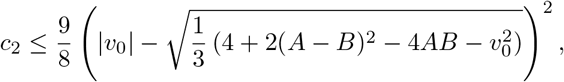

and

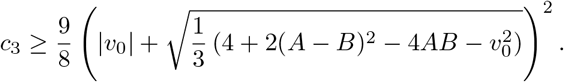

##### Step 3) For small *A*, *B* the intersection *R*(*A, B*) ⋂ (*c*_1_, *c*_2_) is non empty

For this step, let us assume hypotheses i. and ii. detailed in Theorem 2, item (b). We need to find some small *c* > 0 that belongs to every *R_α_*, 0 ≤ *α* ≤ 1. For this purpose we write *R_α_* = {*c* > 0 : *Q*(*c*, *α*) > 0} and we bound

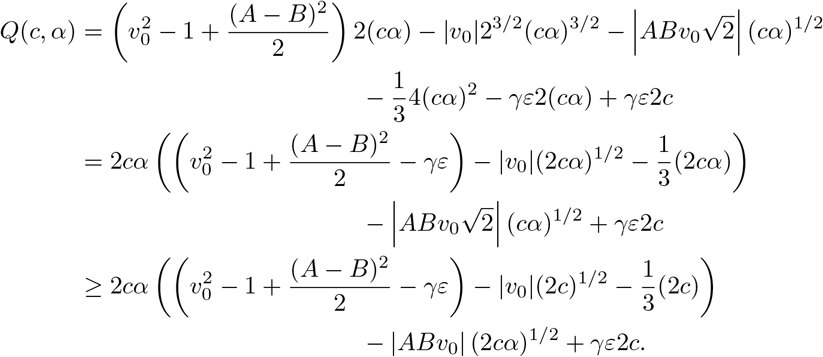

Here we have a quadratic polynomial in 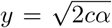 and we can use that 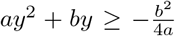 provided that *a* > 0. From hypothesis ii. we know that the coefficient of the quadratic term is positive for *c* > 0 small enough, then we get the following bound

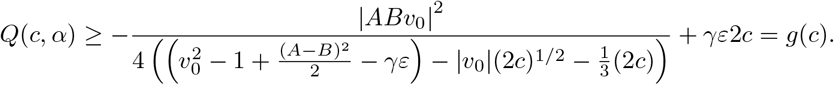

Using ii. again we can take *δ*_0_ = *v*_0_(*A, B*)^2^ –1–*γε* > 0. By continuity of the solution map in (5) near (*A, B*) = (0,0) we can take *K*_1_ > 0 small enough such that |*v*_0_(*A, B*)^2^ – *v*_0_(0,0)^2^| ≤ *δ*_0_/3 provided that max{|*A*|, |*B*|} ≤ *K*_1_. Next, we choose *k* > 0 small enough so that

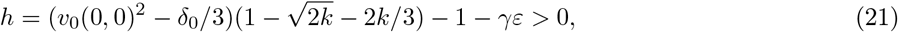

which implies that

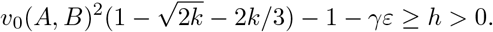

This means that there exists some *K*_0_ ≤ *K*_1_ such that condition i. leads to 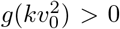. Lastly, we need to verify that 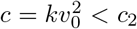. To do this, we notice that due to assumption (6) the bound on *c*_2_ is applicable. Next, since 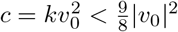, and 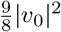 belongs to the forbidden interval obtained in Step 2), we conclude that *c* must in fact be smaller than the lower bound of such interval. Then,

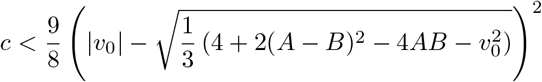

and we conclude that 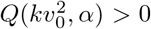 for any *α* ∈ [0,1] and 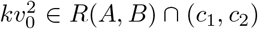. This concludes the proof of item (b) of Theorem 2.

#### Proof of item (c)

To obtain item (c), we use Step 3) of item (b). That proof tells us that given *k* > 0 satisfying (21), there exists some *K*_2_ ≤ *K*_1_ such that if max{|*A*|, |*B*|} ≤ *K*_2_ then *kv*_0_(*A, B*)^2^ ∈ *R*(*A, B*). This means that if we take *δ*_1_ = min{*δ, δ*_0_}, 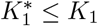 such that |*v*_0_(*A, B*) – *v*_0_(0,0)| ≤ *δ*_1_/2 for 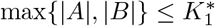 and *k* > 0 small enough such that it satisfy (21) and

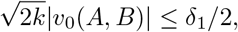

then we get that there exists 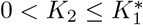 such that *kv*_0_(*A, B*)^2^ ∈ *R*(*A, B*) for max{|*A*|, |*B*|} ≤ *K*_2_. This means that if (*x*_0_, *y*_0_) satisfies

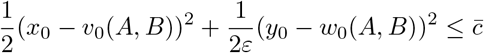

for 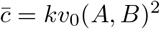 then item (a) implies that

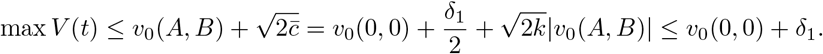

This conclude the proof of Theorem 2.

### 3.2 Proof of Theorem 3

This result shows that, to excite neurons using interferential current stimulation, it is determinant that the beat frequency *η* is not too high. If *η* is high, classical averaging indicates that no neuron activation is possible.

Let (*V, W*) be the solution to system (4). Consider the change of variables *t* → *s/η*, so that we get

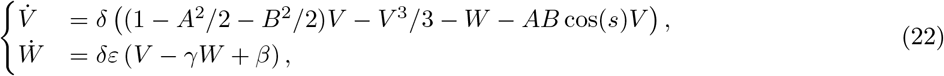

where *δ* = 1/*η*. Since the right-hand side of (22) is periodic in s with period 2π, the averaged system associated to (22) is given by

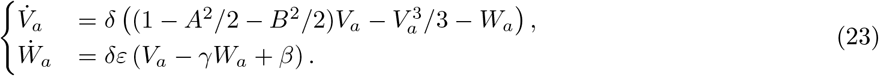

Notice that (*v*_0_, *w*_0_) given by (5) is the equilibrium point of (23) and that the Jacobian matrix *M* of that system at the equilibrium (*v*_0_, *w*_0_) is

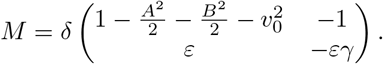

From the Routh-Hurtwitz criterion we get that the equilibrium (*v*_0_, *w*_0_) will be exponentially stable if

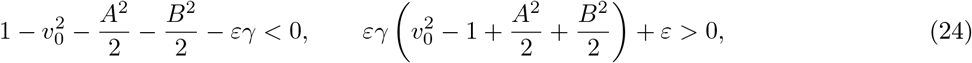

which is always satisfied because condition (2) implies that the solution of (5) is unique and *v*_0_ < –1 for any 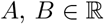. Additionally, it is clear that any sufficiently small ball around (*v*_0_, *w*_0_) is completely contained in the attraction domain of the stable equilibrium. Then, in virtue of the Averaging Theorem [12, Theorem 10.4], [7, Theorem 4.1.1] we get that there exists *δ*\* > 0 such that for all *δ* ≤ *δ*\* the unique solution of (22) is a periodic orbit around (*v*_0_, *w*_0_) that satisfies

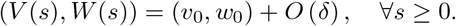

Finally, returning to the *t* variable we conclude that for all 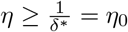 we have

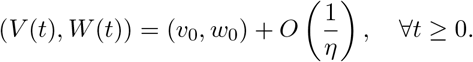

This conclude the proof of Theorem 3.

### 3.3 Properties of the solution of the Partially Averaged System

The last result of this section establishes some properties of the solution for the Partially Averaged System for cases in which no action potentials are elicited. These properties will be used in the proof of the approximation result in Theorem 1.

#### Proposition 6.

*There exist constants M* = *M*(*ε*, *γ*, *β*) > 0 *and c*_0_ = *c*_0_(*ε*, *γ*, *β*) > 0 *such that if* max{|*A*|, |*B*|} ≤ *M and the initial condition* (*x*_0_, *y*_0_) *satisfies*

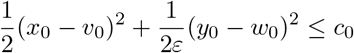

*where* (*v*_0_, *w*_0_) *is given by* (5), *then there exists constants K*_1,1_, *K*_1,2_, *K*_2_ > 0 *such that the solution* (*V, W*) *of system* (4) *satisfies*

i. sup_*t*>0_ *φ*_1,1_(*t*) ≤ –*K*_1,1_ < 0,
ii. sup_t>0_ |*φ*_1,2_(*t*)| ≤ *C* max{*A, B*} = *K*_1,2_,
iii. sup_*t*>0_ |*φ*_2_(*t*)| ≤ *K*_2_,

*where φ*_1,1_(*t*), *φ*_1,2_(*t*), **φ*_2_(*t*) are functions of V*(*t*) *defined by* (12), (13), (14). *Moreover, there exists* 0 < *M*_2_(*ε*, *γ*, *β*) ≤ *M*_1_ *such that if* max{|*A*|, |*B*|} ≤ *M*_2_(*ε*, *γ*, *β*) *then we can guarantee that K*_1,2_ ≤ *K*_1,1_/3.

*Proof*. For item i. we first note that

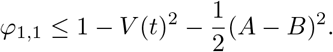

From Theorem 2, we know that if

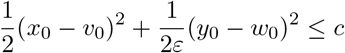

for *c* ∈ *R*(*A, B*) then

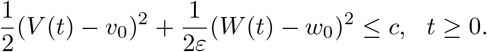

Applying this to the bound of *φ*_1,1_ we get

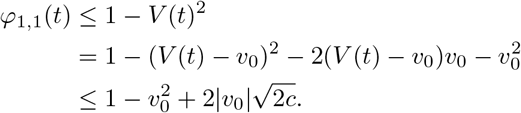

Therefore, since 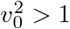, it is enough to choose *c\** > 0 such that

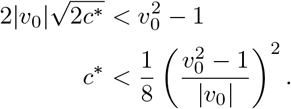

Finally, because of Theorem 2 item iii., given *c\** > 0 there exists *M* = *M*(*ε*, *γ*, *β*, *c\**) > 0 such that if max{|*A*|, |*B*|} ≤ *M* then *c*_0_ ∈ *R*(*A, B*) for some *c*_0_ ≤ *c*\*. Taking 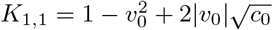 completes the proof of item i.

For item ii., we want to show that we can make *φ*_1,2_ as small as we want by taking max{|*A*|, |*B*|} small. From the definition of *φ*_1,2_, if |*A*| ≤ 1, |*B*| ≤ 1 we can bound

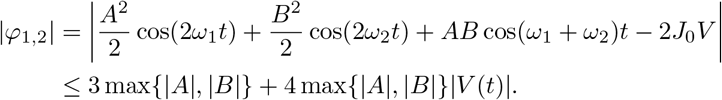

Finally, due to Theorem 2 solutions of system (4) are bounded for all *t* > 0, and we get that there exists some *R* = *R*(*ε*, *γ*, *β*) > 0 such that

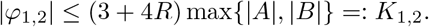

Moreover, by taking 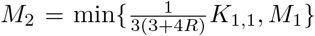, we obtain that *K*_1,2_ ≤ *K*_1,1_/3. This concludes the proof of item ii.

Item *iii*. only requires that the solution stays globally bounded, which we already know by Theorem 2. We get from definition of *φ*_2_

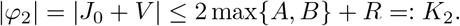

This concludes the proof of Proposition 6.

## 4 Approximation of the FHN system with IFC inputs

In order to study the equation of the approximation error given by Proposition 4, let us consider 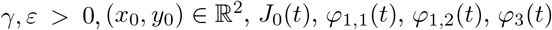 given by (11), (12), (13) and (15), respectively. We define the following linear system

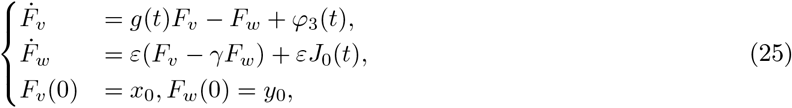

where *g*(*t*) = (*φ*_1,1_(*t*) + *φ*_1,2_(*t*)). Our goal is to study the properties of system (10) for small values of the parameters *A, B*. We perform the analysis in the following three steps

Step 1) Study the stability of the linear system (25).
Step 2) Compare the linear system considered in Step 1) with the equation of the approximation error (10).
Step 3) Conclude that under appropriate conditions the properties of the solution of the Partially Averaged System can be used to conclude that the FHN system stays in a neighborhood of the point (*v*_0_, *w*_0_) given by (5). This is the proof of Theorem 1.

### 4.1 Step 1) Stability of the linear system

We expect that when the error is small in system (10), the linear part is dominant in contrast with the other higher order terms. For that reason, we want to show that the linear system (25) is stable even when a highly oscillatory drift is affecting the system.

#### Proposition 7

(Stability linear system). *Let* (*F_v_, F_w_*) *the solution of system* (25) *and suppose there exists some* 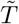 *such that*

i. 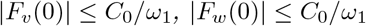,
ii. 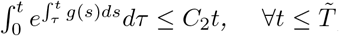,
iii. 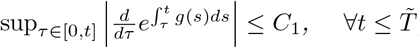,

*for some constants C*_0_, *C*_1_, *C*_2_ > 0. *Then, there exist a constant C*_3_ > 0 *and some* 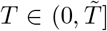 *independent of ω*_1_ *such that*

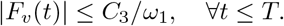

*Proof*. By multiplying the first equation in (25) by 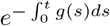 we get

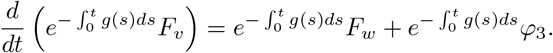

We integrate in [0, *t*] to obtain

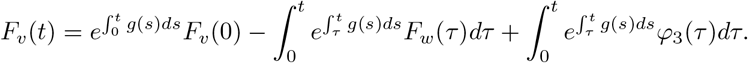

Analogously, multiplying the second equation in (25) by *e^εγt^* and integrating in [0, *t*], we get

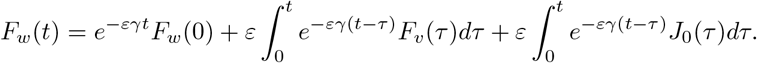

Next, we consider the following Picard’s iteration. Set 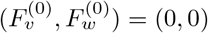 and for *k* > 1

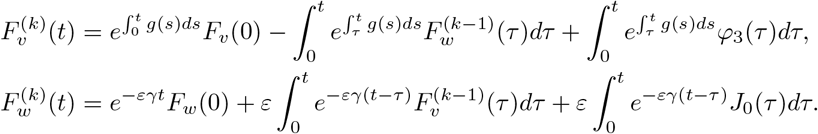

To study the convergence we look at the difference between two consecutive steps

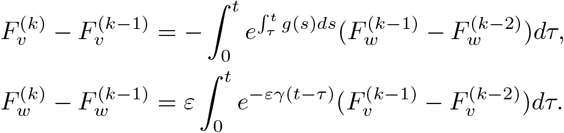

Next, for 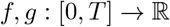 we consider the norm

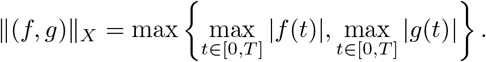

Then, we can write the estimate

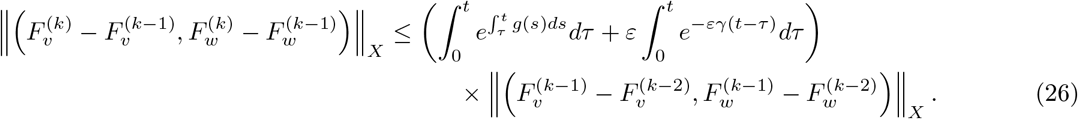

Now, we choose 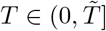 so that 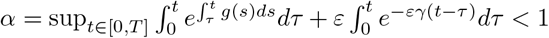. Then, the solution can be written as

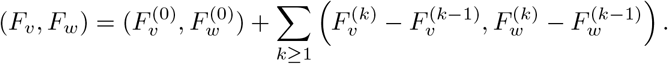

Applying estimate (26) we get

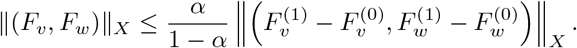

Now, we estimate the first Picard’s iteration. Let us consider the following

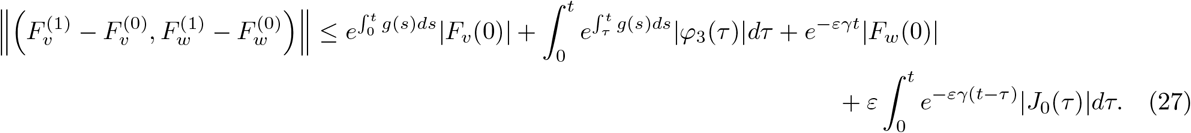

In order to get appropriate bounds for (27) we need the following lemma whose proof can be found later in the Appendix A.

#### Lemma 8

(Integral estimate for slow varying functions). *Let f* ∈ *C*^1^([0, *T*]) *and ω* ≫ 1 *then*

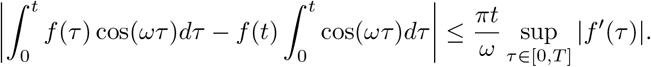

We apply Lemma 8 to get suitable bounds for

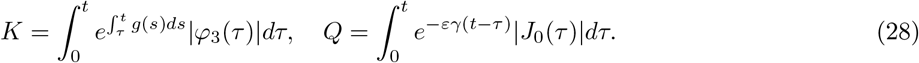

Let us begin by *K*. Recalling the definition of *J*_0_ and *φ*_3_ we have

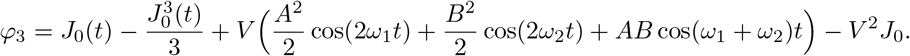

We can re-write *φ*_3_(*t*) as follows,

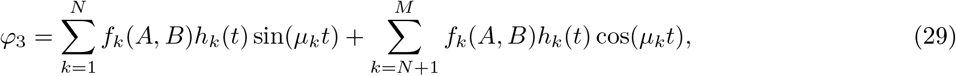

where 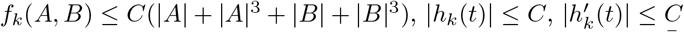 and *μ_k_* ≥ *ω*_1_/2. Then, in virtue of Lemma 8 and hypothesis iii., for each *k* = 1, ⋯, *N* there exists a constant 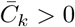 such that

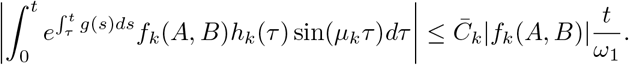

Analogously, for each *k* = *N* +1, ⋯, *M* there exists a constant 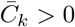 such that

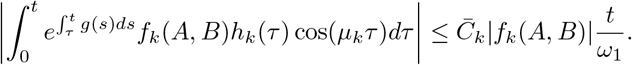

Let us define 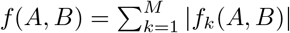. Finally, we get that

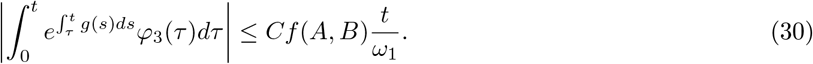

for some constant *C* > 0. Now, for *Q* we apply Lemma 8 to get that

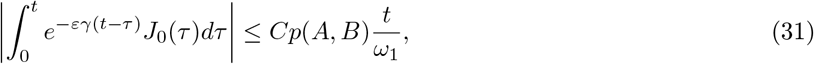

where *f*(*A, B*), *p*(*A, B*) ≤ *C*(|*A*| + |*B*| + |*A*|^3^ + |*B*|^3^) and *C* > 0. Applying (30) and (31) to (27) we get

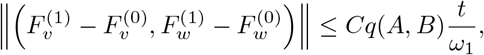

for a function 0 ≤ *q*(*A, B*) ≤ *C*(|*A*| + |*B*| + |*A*|^3^ + |*B*|^3^) and a constant *C* > 0. Lastly, we need to confirm that the size required for *α* is not too big. For this purpose we notice that the condition *α* < 1 give us

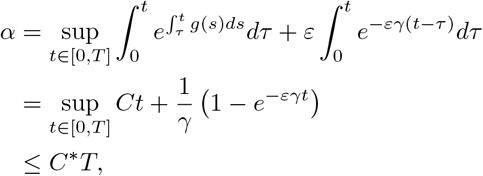

by hypothesis ii. Then, we conclude that 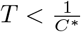 works. This concludes the proof of Proposition 7.

**Remark 4.1.** *To check the hypothesis ii. in Proposition 7 we can notice that because of our choice of g*(*t*) *we know that* 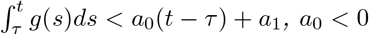, *which imply hypothesis ii*.

### 4.2 Step 2) Comparison with linear system

The following step, in the proof of the approximation estimate is to show that the linear estimate provided by Proposition 7 is still valid for the full problem under appropriate conditions.

#### Proposition 9

(Comparison with the linear problem for system (10)). *Let* 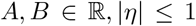, *fix ε*, *γ*, *β* > 0 *satisfying* (2). *Consider the solution* (*E_v_, E_w_*) *of system* (10) *and let* (*F_v_, F_w_*) *be the solution to the linear system* (25). *There exists M*(*ε*, *γ*, *β*) > 0 *such that if* max{|*A*|, |*B*|} ≤ *M*(*ε*, *γ*, *β*) *and for some* 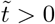 *the solution* (*V, W*) *of the Partially Averaged System* (4) *satisfies for* 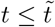 *that*

i. *φ*_1,1_(*t*) ≤ –*K*_1,1_ < 0,
ii. |*φ*_1,2_ (*t*) | ≤ *K*_1,2_,
iii. |*φ*_2_(*t*)| ≤ *K*_2_,

*where φ*_1,1_(*t*), *φ*_1,2_(*t*), *φ*_2_(*t*) *are functions of V*(*t*) *defined by* (12), (13), (14), *and K*_1,2_ ≤ *K*_1,1_/3. *Then, there exists M*_2_ > 0 *such that if* 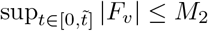, *then difference between both systems* (*R_v_, R_w_*) = (*E_v_* – *F_v_*, *E_w_* – *F_w_*) *satisfies*

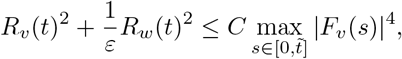

*for* 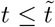 *and a constant* 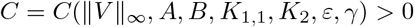.

*Proof*. Let us consider

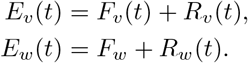

The system equation for (*R_v_*(*t*), *R*_2_(*t*)) is given by

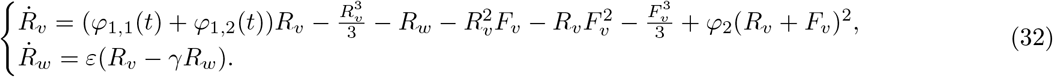

Multiplying the first equation by *R_v_*, the second one by 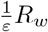, and adding them together we get

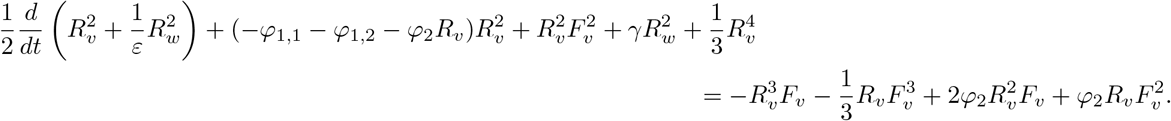

Using Young’s inequality we obtain the following bounds. For all *ϵ* > 0, *δ* > 0 and *δ*_2_ > 0, it holds

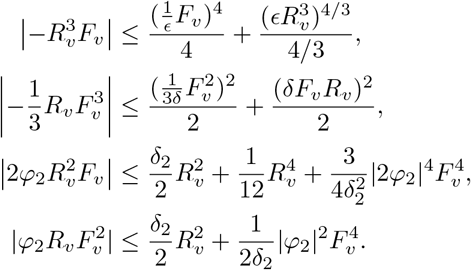

Next, by assumptions i., ii. and iii. we can choose *ϵ, δ* > 0 such that 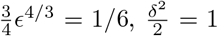 and *δ*_2_ = *K*_1,1_/3. Thus we get

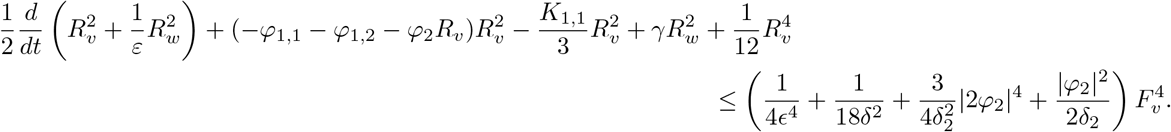

Recalling again assumptions i., ii., and iii. we obtain

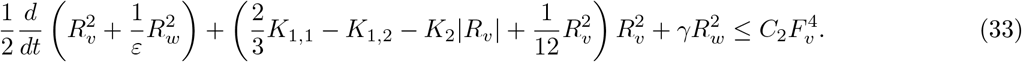

Next, by the assumption 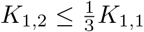, we can write

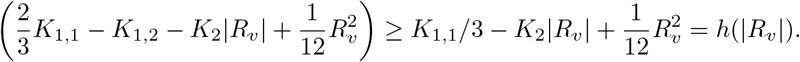

Here we have two cases, if the discriminant 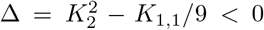 then the polynomial is always positive. Therefore, there exists *c*_0_ = *c*_0_(*K*_1,1_, *K*_2_) > 0 such that

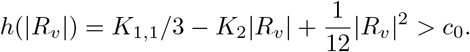

On the other hand, when 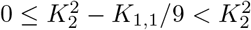 we know that if we denote by 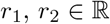 the roots of *h*(*x*) = 0 then *r*_1_ and *r*_2_ have the same sign and therefore *h*(*x*) is monotone in [0, min{|*r*_1_|, |*r*_2_|}]. This implies that taking

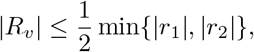

we can find a constant *c*_0_ = *c*_0_(*K*_1,1_, *K*_2_) > 0 such that

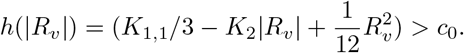

We obtain that whenever 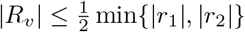 we get from (33) that

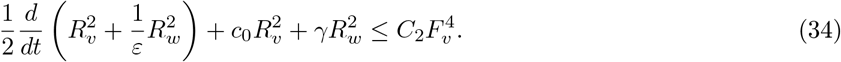

Defining *c*_3_ = min{*c*_0_, *γε*}, we see that (34) becomes

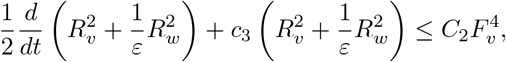

and by considering the integral factor *e*^2*c*_3_*t*^, we get

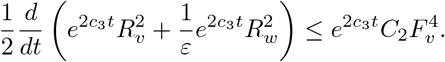

For 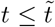, we can integrate and use that *R_v_* (0) = *R_w_* (0) = 0 to get

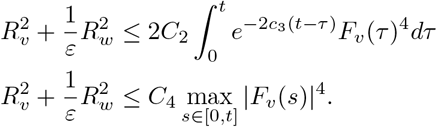

In the case where 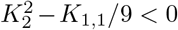 we are done, if not, we have to check that the condition 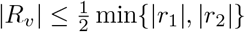 is preserved. For this we we require that

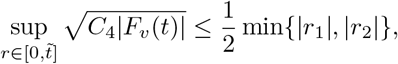

which is satisfied for *M*_2_ > 0 small enough. This concludes the proof of Proposition 9.

**Remark 4.2.** *We are interested on this approximation because from the stability of the linear system* (25) *we expect that F_v_*(*t*) = *O*(1/*ω*_1_) *for some time t* ≤ *T*. *This result indicates that whenever this condition is true, the nonlinear error system* (10) *also satisfies such estimates for the same time range and for large enough frequencies*.

### 4.3 Step 3) Proof of Theorem 1

Let *c*_0_ > 0 be as in Proposition (6). Take the initial conditions for the the Partially Averaged System to be the same as for the FHN system. Then by assumption we have

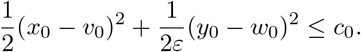

Next, by Proposition 6 we know that the solution (*V, W*) of the Partially Averaged System satisfies the hypothesis of Proposition 7. This means that there exists 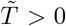 independent of *ω*_1_ such that the solution (*F_v_, F_w_*) of (25) satisfies 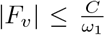 for 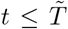. And by taking *ω*_1_ ≥ *ω*_0_ = *ω*_0_(||*V*||_∞_, *A, B, K*_1,1_, *K*_2_, *ε*, *γ*) we can guarantee that 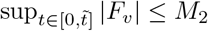 and therefore the hypotheses of Proposition 9 are satisfied. As well, because of the definition of *E_v_*, *E_w_* in Proposition 4, we conclude that

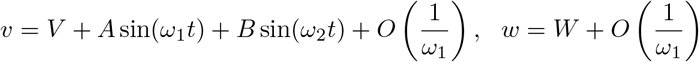

for 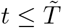. This concludes the proof of Theorem 1.

## 5 Simulations

To illustrate the behavior of the Partially Averaged System introduced here and its usefulness to study the full FHN system under IFC stimulation, we performed three simple experiments in which we measured neuron responses to IFC stimulation. All the simulations were performed using Mathematica 12.3, and the source codes are available at https://github.com/estebanpaduro/fhn-pas-simulations. In all simulations we set *ε* = 0.08, *β* = 0.8, *γ* = 0.5. Additionally, according to our parameter selection, we set a threshold of *v* > 1 for action potential identification, as explained in Subsection 1.4.

The first experiment illustrates the averaging process (Figure 3). In this simulation, we used *A* = 0.5, *B* = 0.5, 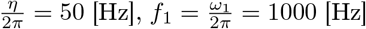, and a simulation time of 100 [ms]. Figure 3a shows the solution of system (1). In this case, three action potentials were generated, and a high frequency signal overlapped. Figure 3b shows the solution of the FHN system after removing the high frequency signal used in the proposed decomposition of the solution (*v, w*) to the FHN system (1), given by (8), to obtain the system for which the averaging will be applied. Finally, Figure 3c shows the solution of the Partial Averaged System (4), which is non-autonomous and has an explicit dependence on the beat frequency, *i.e*. frequency of the envelope, generated by the interference of the KHF input signals.

**Figure 3:**
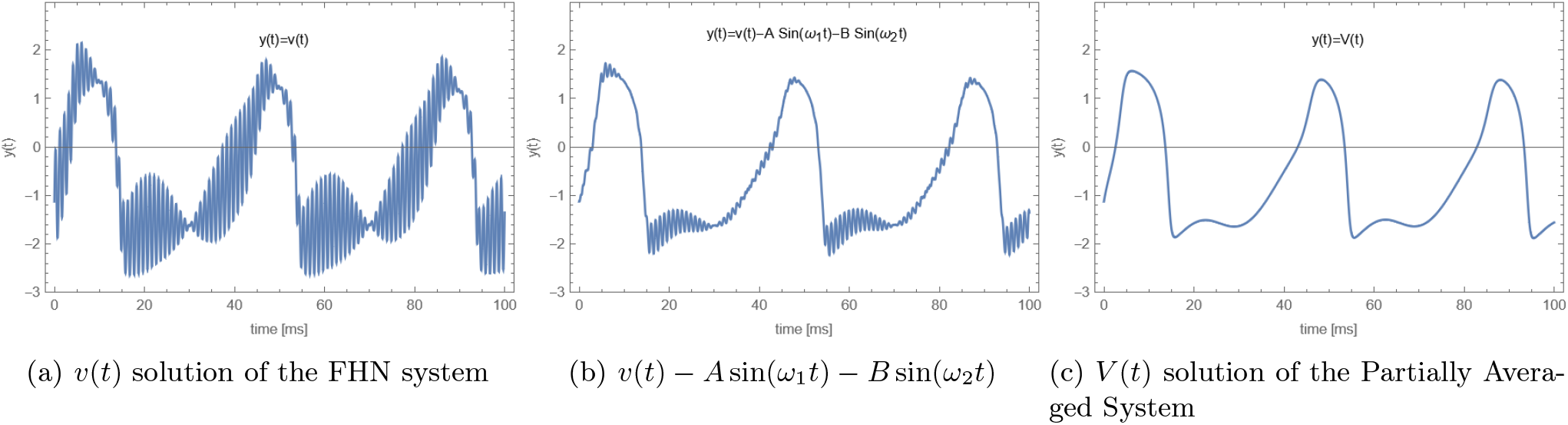
Partial Averaging for the FHN system. The dynamics of the membrane voltage, *v*(*t*), is shown for the full FHN system and the Partially Averaged System. Three action potentials or spikes were generated in this case

In the second experiment, we varied the parameters *A* and *B* in the range 0–1.5 with a step size of Δ = 0.02, a frequency *η*/(2π) of 50 [Hz], and a simulation time of 1 [s]. As model output, we measured the spike rate, *i.e*. number of action potentials of the model neuron per unit time, and we examined the most effective parameter combination for neuron activation. Figure 4 shows the spike rate for four different values of *η*. In this case, since *A* and *B* represent the magnitude of each sinusoid, respectively, they indirectly define the interference region. The results of this simulation suggest that the interference field is rather small, and its size depends on the beat frequency *η*. Further, there is an optimal beat frequency that maximizes the spike rate in about 30 [spikes/s], a result further confirmed in the third experiment.

**Figure 4:**
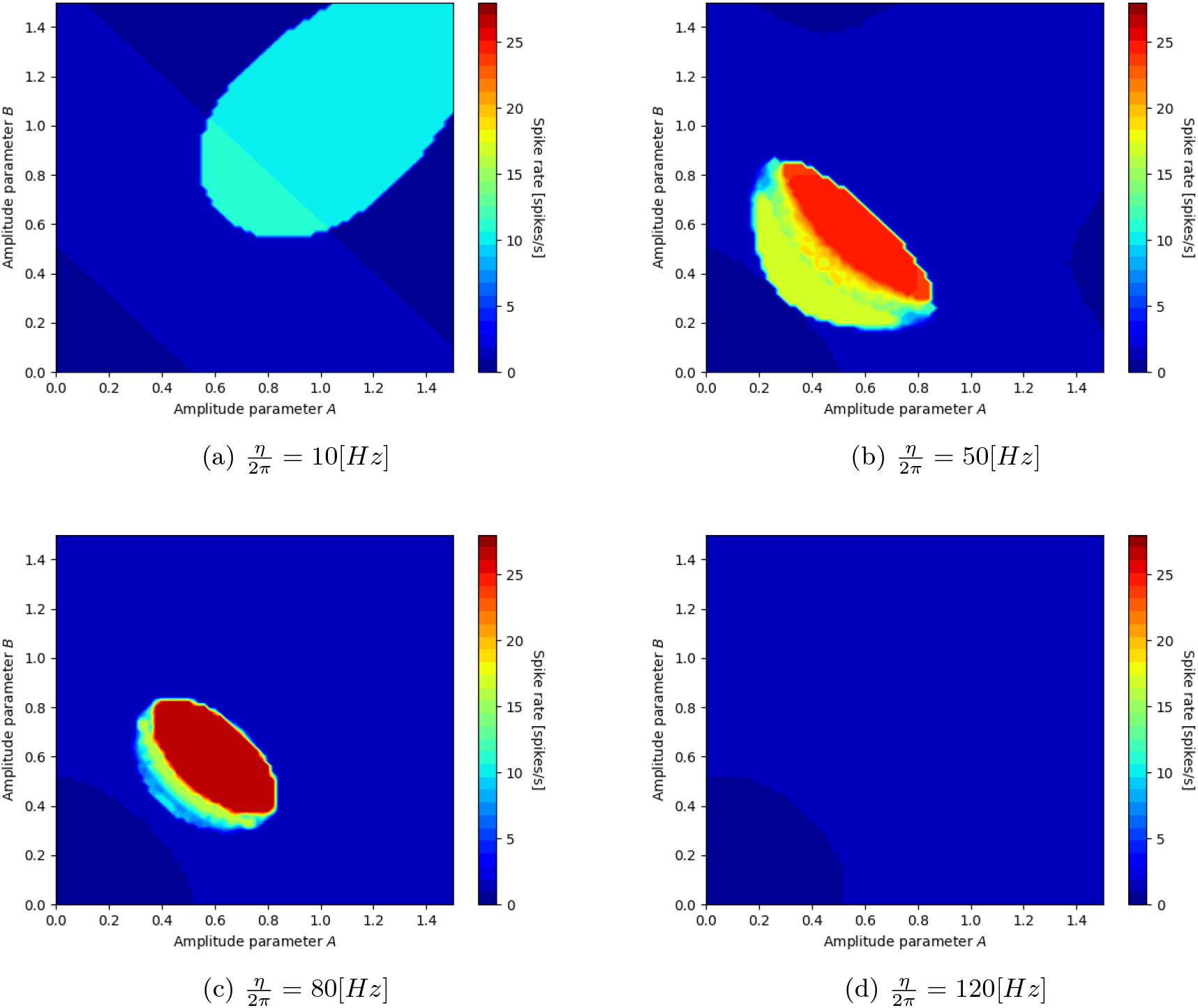
Spike rate –number of action potential per unit time– for the Partially Averaged System (4) as a function of the amplitude, *A* and *B*, of the stimulation signals. The color plots represent a beat frequency, *η*, of 10, 50, 80 and 120 [Hz], respectively

The third experiment considered the effects of varying the beat frequency *η*. Here, we set the amplitude of both signals to the same magnitude, which we varied in the range 0–3 with a step size of 0.01. For the beat frequency, we varied *η*/(2π) in the range 0–200 [Hz] with a step size of 1 [Hz]. In this case, the simulation time was of 1 [s], and we measured the number of action potentials elicited in that interval. Figure 5 shows that only for beat frequencies lower than ~ 100 [Hz] it is possible to activate neurons, and within a narrow amplitude range.

**Figure 5:**
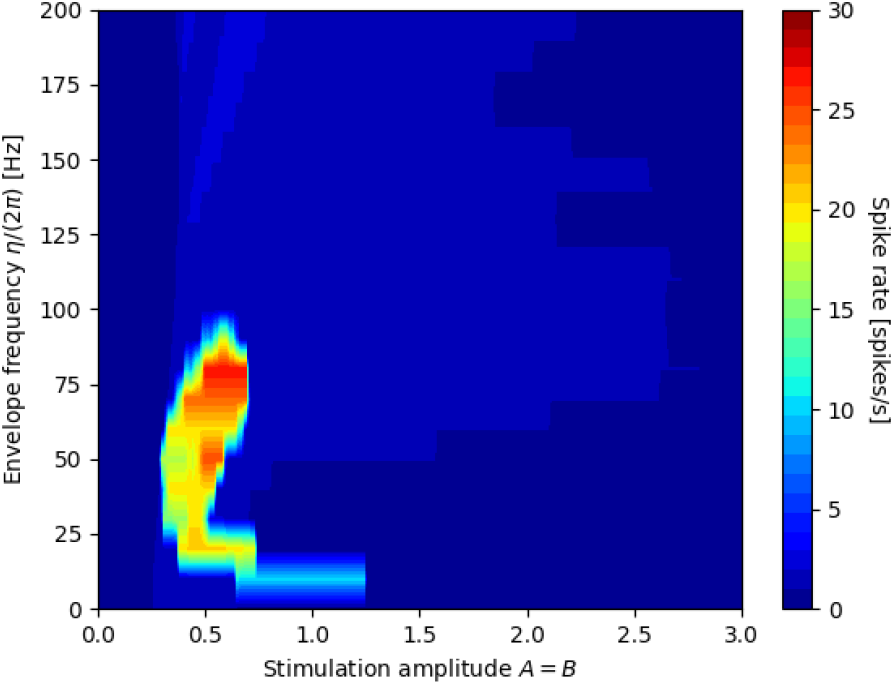
Spike rate as a function of the beat frequency, *η*, and the amplitude of the stimulation signal for the Partially Averaged System (4). In this case, *A* = *B*

## 6 Conclusions

In this study, we introduced a Partially Averaged System to model a FHN neuron subjected to IFC stimulation. Our system is computationally inexpensive, and shows explicit dependence of the neuron variables on the IFC parameters, namely, the beat frequency and the amplitude of each KHF input signal. We determined conditions under which this system is a good approximation of the complete FHN system, and presented some properties that can be used to predict neuron responses to IFC stimulation. From a mathematical viewpoint the biggest challenge was to deal with non-autonomous systems and obtain useful estimates allowing us to state exploitable results. Future work may use similar averaging approaches to analyze partial derivative systems with spatial variables, *e.g*., the non-space-clamped FHN nerve model [19], or stochastic FHN approaches [25]. This would allow a more comprehensive analysis of the effects of IFC on morphologically detailed model neurons.

Our numerical results suggest that the intended interference effect is limited to a narrow range of stimulation parameters, so that for neuron activation using surface electrodes, fine adjustment of the stimulation settings may be necessary. Further, a precise control of signal modulation within the interference field may be required as the beat frequency appears to strongly determine neuron activation. Our novel description of the IFC technique contributes to the understanding of the mechanisms of neurostimulation using this type of signals, and can have implications in the design of more efficient and/or effective noninvasive therapies.

## A Appendix: Technical proofs

### A.1 Proof of Proposition 4

Denote *E_v_* = *v* – *V* – *J*_0_(*t*), *E_w_* = *w* – *W* with *J*_0_(*t*) = *A* sin(*ω*_1_*t*) + *B* sin(*ω*_2_*t*), notice that 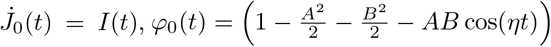. Using 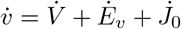 we can combine equations (1) and (4) to obtain

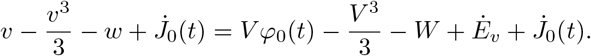

Substituting *v* = *V* + *E_v_* + *J*_0_, *w* = *W* + *E_w_* we get

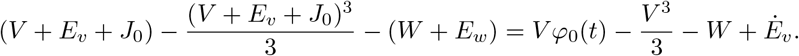

Gathering terms together we get the following equation for *E_v_*

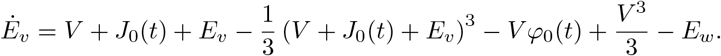

By expanding the cubic term, this equation can be written as

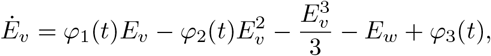

where the functions 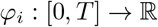, *i* = 1, 2, 3 are given by

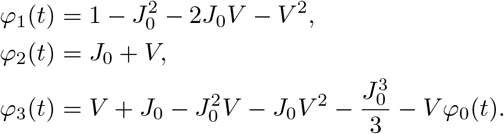

To obtain an equation for *E_w_* we substitute *v* = *V* + *E_v_* + *J*_0_, *w* = *W* + *E_w_* in equation (1) to obtain

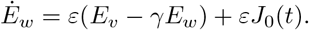

Next, we consider the following decomposition for *φ*_1_

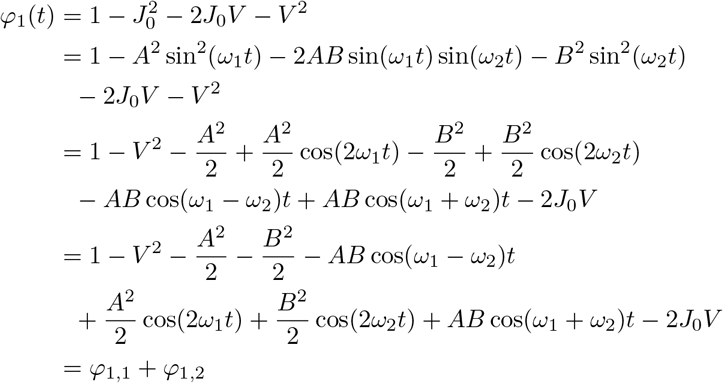

where

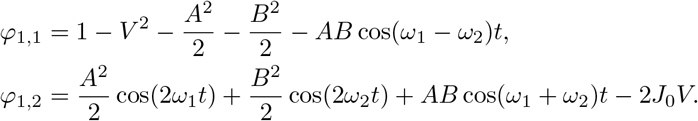

The idea is that *φ*_1,1_ gathers the slow varying terms and *φ*_1,2_ the highly oscillatory terms. For the term *φ*_3_(*t*) we need the following

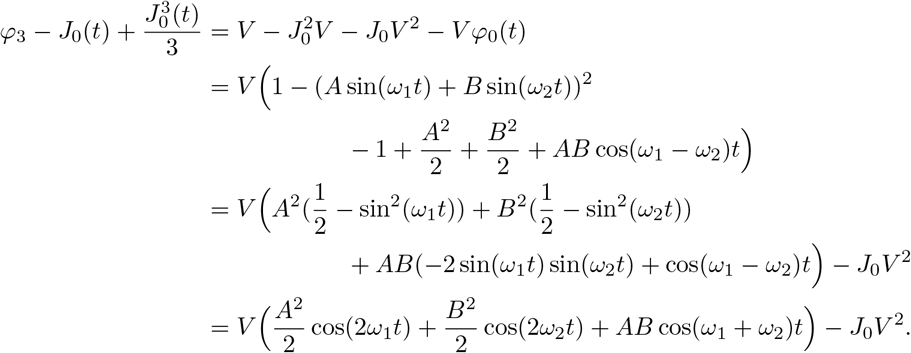

Thus, we can write

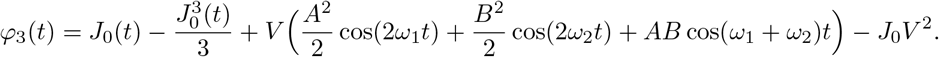

Putting all together we obtain the equations for (*E_v_, E_w_*)

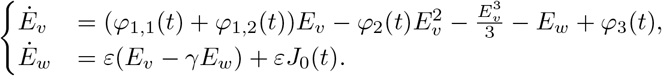

### A.2 Proof of Lemma 5

We argue by contradiction, if this is not the case, then given *δ*_0_ > 0 there exists *t*_0_ ∈ (0, ∞) and 0 < *δ*_1_ ≤ *δ*_0_ such that *f*(*t*_0_) = *c* and *f*(*t*) > *c* for *t* ∈ (*t*_0_, *t*_0_ + *δ*_1_). Now because 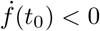, by continuity there exists some 0 < *δ*_2_ < *δ*_1_ such that 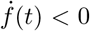 for *t* ∈ [*t*_0_, *t*_0_ + *δ*_2_]. Next, by the mean value theorem there exists some *z* ∈ (*t*_0_, *t*_0_ + *δ*_2_) such that

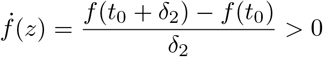

but this is a contradiction without choice of *δ*_2_. Therefore, we conclude f (*t*) ≤ *c* for all *t* ∈ (0, ∞).

### A.3 Proof of Lemma 8

The key for the estimate is to split the interval in full periods of the function cos(*ωt*). Let *N* = ⌊*t*/(2π/*ω*)⌋ then for *k* = 0, ⋯, *N* – 1 we consider the estimate

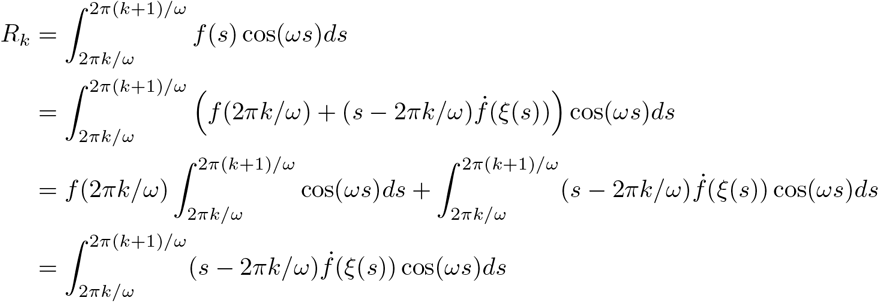

Thus, we get the bound

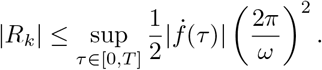

The estimate is different for the last segment of the interval, in fact,

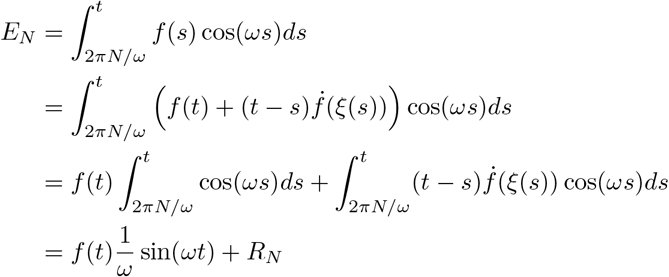

and the estimate is

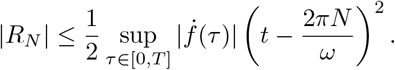

Finally, we can combine the estimates to obtain

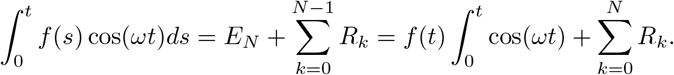

We get in this way that

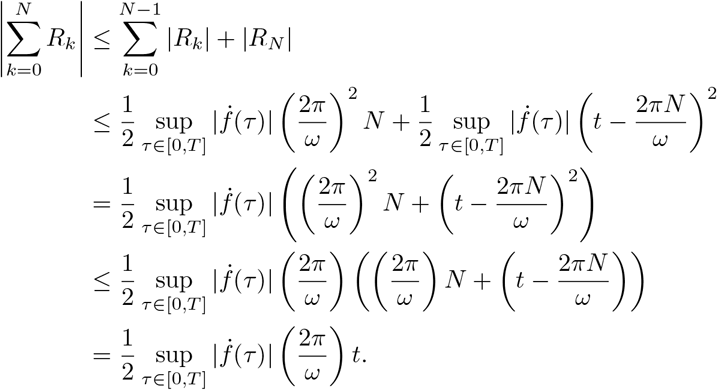

This concludes the proof of Lemma 8.

## Notes

* This work has been partially supported by ANID Millennium Science Initiative Program trough Millennium Nucleus for Applied Control and Inverse Problems NCN19-161, Basal Project FB0008 AC3E, and Fondecyt 11190822.

### Competing Interest Statement

The authors have declared no competing interest.

